# A pedagogical walkthrough of computational modeling and simulation of Wnt signaling pathway using static causal models in Matlab

**DOI:** 10.1101/011064

**Authors:** Shriprakash Sinha

**Author notes:** Shriprakash Sinha drafted the manuscript. E-mail: Shriprakash Sinha drafted the manuscript.

## Abstract

**Abstract:** A tutorial introduction to computational modeling of Wnt signaling pathway in a human colorectal cancer dataset using static Bayesian network models is provided. The walkthrough might aid bio-logists/informaticians in understanding the design of computational experiments that is interleaved with exposition of the Matlab code and causal models from Bayesian Network toolbox. This is done in order to ease the understanding of beginner students and researchers in transition to computational signaling biology, who intend to work in the field of modeling of signaling pathways. The manuscript expounds the computational flow of the contents in advance article^1^ via code development and takes the reader in a step by step process of how • the collection and the transformation of the available biological information from literature is done, • the integration of the heterogeneous data and prior biological knowledge in the network is achieved, • conditional probability tables for nodes in biologically inspired tables are estimated, • the simulation study is designed, • the hypothesis regarding a biological phenomena is transformed into computational framework, and • results and inferences drawn using *d*-connectivity/separability are reported. The manuscript finally ends with a programming assignment to help the readers get hands on experience of a perturbation project. Matlab code with dataset is made available under GNU GPL v3 license at google code project on https://code.google.com/p/static-bn-for-wnt-signaling-pathway

**Insight, Innovation and Integration:** Simulation study involving computational experiments dealing with Wnt signaling pathways abound in literature but often lack a pedagogical perspective that might ease the understanding of beginner students and researchers in transition who intend to work on modeling of the pathway. This paucity might happen due to restrictive policies which enforce an unwanted embargo on the sharing of important scientific knowledge. The manuscript elucidates embedding of prior biological knowledge, integration of heterogeneous information, transformation of biological hypothesis into computational framework and design of experiments in a simple manner interleaved with aspects of Bayesian Network toolbox and Matlab code so as to help readers get a feel of a project related to modeling of the pathway.

## 1 A journey of thousand miles begins with a single step

A tutorial introduction to computational modeling of Wnt signaling pathway in a human colorectal cancer dataset using static Bayesian network models is provided. This work endeavours to expound in detail the simulation study in Matlab along with the code while explaining the concepts related to Bayesian networks. This is done in order to ease the understanding of beginner students and researchers in transition to computational signaling biology, who intend to work in the field of modeling of signaling pathways. The manuscript elucidates embedding of prior biological knowledge, integration of heterogeneous information, transformation of biological hypothesis into computational framework and design of experiments in a simple manner interleaved with aspects of Bayesian Network toolbox and Matlab code so as to help readers get a feel of a project related to modeling of the pathway. Programming along with the exposition in the manuscript could clear up issues faced during the execution of the project.

This manuscript uses the contents of the advance article Sinha^1^ as a basis to explain the workflow of a computational simulation project involving Wnt signaling pathway in human colorectal cancer. The aim of Sinha^1^ was to computationally test whether the activation of *β*-*catenin* and *TCF*4 based transcription complex always corresponds to the tumorous state of the test sample or not. To achieve this the gene expression data provided by Jiang *et al.*^3^ was used in the computational experiments. Further, to refine the model, prior biological knowledge related to the intra/extracellular factors of the pathway (available in literature) was integrated along with epigenetic information.

Theory and programming code will be explained in an interleaved manner to help the readers get an insight into the hurdles faced will executing such a project. Material from Sinha^1^ will be presented in grey colored boxes and used to explain the various aspects of the Matlab code presented here. Code will be presented in `typewriter` font and functions in the text will be presented in sans serif.

## 2 Modeling and simulation

### 2.1 Data collection and estimation

An important component of this project is the Bayesian Network Toolbox provided by Murphy *et al.*^15^ and made freely available for download on https://code.google.com/p/bnt/ as well as a Matlab license. Instructions for installations are provided on the website. One can make a directory titled *temp* with a subdirectory named *data* and transfer the *geneExpression.mat* file into *data*.

~~~
>> mkdir temp
>> cd temp
>> mkdir data
>>
~~~

The .mat file contains expression profiles from Jiang *et al*.^3^ for genes that play a role in Wnt signaling pathway at an intra/extracellular level and are known to have inhibitory affect on the Wnt pathway due to epigenetic factors. For each of the 24 normal mucosa and 24 human colorectal tumor cases, gene expression values were recorded for 14 genes belonging to the family of *SFRP*, *DKK*, *WIF*1 and *DACT*. Also, expression values of established Wnt pathway target genes like *LEF*1, *MY C*, *CD*44 and *CCND*1 were recorded per sample.

The directory *temp* also contains some of the .m files, parts of contents of which will be explained in the order of execution of the project. The main code begins with a script titled *twoHoldOutExp.m*. This script contains the function twoHoldOutExp which takes two arguments named `eviDence` and `model`. `eviDence` implies the evidence regarding ’ge’ for gene evidence, ’me’ for methylation, ’ge+me’ for both gene and methylation, while `model` implies the network model that will be used for simulation. Sinha^1^ uses three different models i.e ’t1’ or 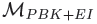 that contains prior biological knowledge as well as epigenetic information, ’t2’ or 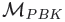 that contains only prior biological knowledge and finally, ’p1’ or 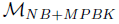 that is a modified version of naive bayes framework from Verhaegh *et al.*^2^. In Matlab, one can type the following

~~~
>> twoHoldOutExp (“ge”, “t1”)
~~~

The code begins with the extraction of data from the gene expression matrix by reading the *geneExpression.mat* file via the function readCustomFile in the *readCustomFile.m* and generates the following variables as the output - (1) `uniqueGenes` - name of genes gleaned from the file, (2) `expressionMatrix` - 2D matrix containing the gene expression per sample data (3) `noGenes` - total number of genes available (4) `noSamples` - total number of samples available (5) `groundTruthLabels` - original labels available from the files (6) `transGroundTruthLabels` - labels transformed into numerals.

~~~
% Data Collection
%=====
% Extract data from the gene expression
% matrix
[uniqueGenes, expressionMatrix,…
noGenes, noSamples, groundTruthLabels,…
transGroundTruthLabels] = …
readCustomFile (’data/geneExpression.mat’);
~~~

### 2.2 Assumed and estimated probabilities from literature

Next, the probability values for some of the nodes in the network is loaded, depending on the type of the network. Why these assumed and estimated probabilities have been addressed in the beginning of the computation experiment will be explained later. Meanwhile, the estimation of probabilities is achieved through the function called dataStorage in the *dataStorage.m*. The function takes the name of the `model` as an input argument and returns the name of the file called *probabilities.mat* in the variable `filename`. The mat file contains all the assumed and computed probabilities of nodes for which data is available and is loaded into the workspace of the Matlab for further use.

~~~
% Load probability values for some of
% the nodes in the network
fname = dataStorage (model);
load (fname);
~~~

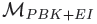 (model=’t1’) requires more estimations that 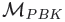 (model=’t2’) and 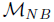 (model=p1) due to use of epigenetic information. Depending on the type of model parameter fed to the function dataStorage the probabilities for the following factors are estimated

1. Repressive Histone Mark *H*3*K*27*me*3 for *DACT*3 11 Loci from Jiang *et al.*^3^ were adopted. Via fold enrichment, the affects of the *H*3*K*27*me*3 was found 500 bp downstream of and near the *DACT*3 transcription start site (TSS) in HT29 cells. These marks were recorded via chromatin immuno-precipitation (ChiP) assays and enriched at 11 different loci in the 3.5 kb to 3.5 kb region of the DACT3 TSS. Fold enrichment measurements of *H*3*K*27*me*3 for normal *FHs*74*Int* and cancerous *SW* 480 were recorded and normalized. The final probabilities are the average of the normalized values.
2. Active Histone Mark *H*3*K*4*me*3 for *DACT*3 Loci from Jiang *et al.*^3^ were adopted. Via fold enrichment, the affects of the *H*3*K me*3 was found 500 bp downstream of and near the *DACT*3 transcription start site (TSS) in HT29 cells. These marks were recorded via chromatin immuno-precipitation (ChiP) assays and enriched at 11 different loci in the 3.5 kb to 3.5 kb region of the *DACT* 3 TSS. Fold enrichment measurements of *H*3*K*4*me*3 for normal *FHs*74*Int* and cancerous *SW* 480 were recorded and normalized. The final probabilities are the average of the normalized values.
3. Fractions for methylation of *DKK*1 and *WIF*1 gene taken from Aguilera *et al.*^16^ via manual counting through visual inspection of intensity levels from methylation specific PCR (MSP) analysis of gene promoter region and later normalized
4. Fractions for methylation and non-methylation status of *SF RP*1, *SF RP*2, *SF RP*4 and *SF RP*5 (CpG islands around the first exons) was recorded from 6 affected individuals each having both primary CRC tissues and normal colon mucosa from Suzuki *et al.*^17^ via manual counting through visual inspection of intensity levels from methylation specific PCR (MSP) analysis of gene promoter region and later normalized.
5. Methylation of *DACT*1 (+52 to +375 BGS) and *DACT*2 (+52 to +375 BGS) in promoter region for *Normal*, *HT*29 and *RKO* cell lines from Jiang *et al.*^3^ was recorded via counting through visual inspection of open or closed circles indicating methylation status estimated from bisulfite sequencing analysis and later normalized.
6. Concentration of *DV L*2 decreases with expression of *DACT*3 and vice versa Jiang *et al.*^3^. Due to lack of exact proportions the probability values were assumed.
7. Concentration of *β*-catenin given concentrations of *DV L*2 and *DACT*1 varies and for static model it is tough to assign probability values. High *DV L*2 concentration or suppression (expression) of *DACT*1 leads to increase in concentration of *β*-catenin (^3^, Yuan *et al.*^18^). Wet lab experimental evaluations might reveal the factual proportions.
8. Similarly, the concentrations of *TRCMP LX* (Clevers^4^, Kriegl *et al.*^19^) and *TCF*4 (Verhaegh *et al.*^2^) have been assumed based on their known roles in the Wnt pathway. Actual proportions require further wet lab tests.
9. Finally, the probability of *Sample* being tumorous or normal is a chance level as it contains equal amount of cancerous and normal cases.

Note that all these probabilities have been recorded in table 1 of Sinha^1^ and their values stored in the *probabilities.mat* file. Addressing the question of why these probabilities have been estimated earlier, it can be seen that the extra/intracellular factors affecting the Wnt pathway in the data set provided by Jiang *et al.*^3^ contains some genes whose expression is influenced by epigenetic factors mentioned in table 2. Hence it is important to tabulate and store prior probability values for known biological factors that influence the pathway. Also, the probability values of these nodes have been computed earlier due to prior available information. Once estimated or assumed based on biological knowledge, these probabilities needed not be recomputed and are thus stored in proper format at the beginning of the computational experiment.

**Table 1.**
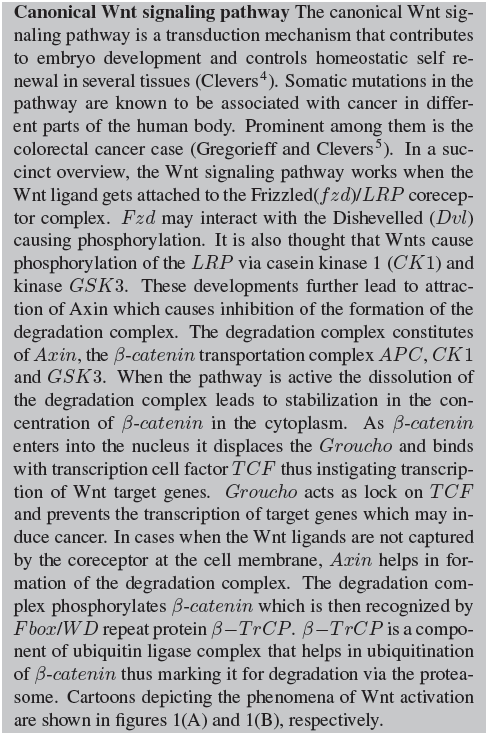
Canonical Wnt Pathway from Sinha^1^

**Table 2.**
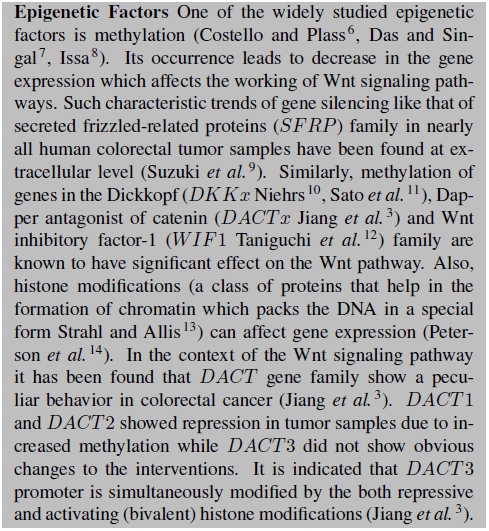
Epigenetic Factors from Sinha^1^

### 2.3 Building the bayesian network model

Next comes the topology of the network using prior biological knowledge made available from results of wet lab experiments documented in literature. This is achieved using the function generateInteraction in the file `generateInteraction.m`. The function takes in the set of `uniqueGenes` and the type of `model` and generates a cell of `interaction` for the Bayesian network as well as a cell of unique set of `Nodenames`. `interaction` contains all the prior established biological knowledge that caries causal semantics in the form of arcs between parent and child nodes. It should be noted that even though the model is not complete due to its static nature, it has the ability to encode prior causal relationships and has the potential for further refinement.

~~~
% Building the Bayesian Network model
%=====
% Generate directionality between
% parent and child nodes
[interaction, nodeNames] = …
 generateInteraction (uniqueGenes,…
 model);
~~~

The `interaction` and `nodeNames` go as input arguments to the function mk_adj_mat, which generates an adjacency matrix for a directed acyclic graph (DAG) stored in `dag`. Using functions biograph and input arguments `dag` and `nodeNames` generates a structure `gObj` that can be used to view the topology of the network. A crude representation of 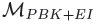 and 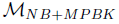 shown in figures 2 and 3 was generated using the function view.

**Fig. 1.**
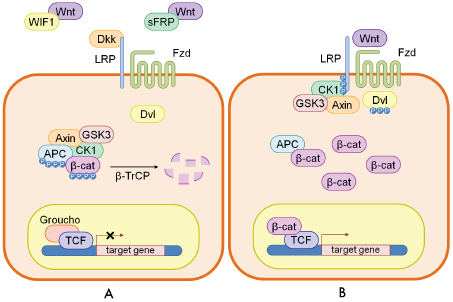
A cartoon of wnt signaling pathway contributed by Verhaegh *et al.*^2^. Part (A) represents the destruction of *β*-*catenin* leading to the inactivation of the wnt target gene. Part (B) represents activation of wnt target gene.

**Fig. 2.**
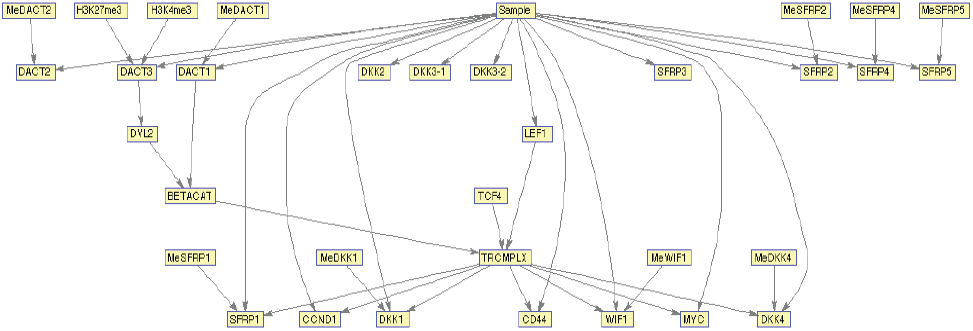
Influence diagram of 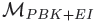 contains partial prior biological knowledge and epigenetic information in the form of methylation and histone modification. In this model the state of Sample is distinguished from state of *TRCMP LX* that constitutes the Wnt pathway.

**Fig. 3.**
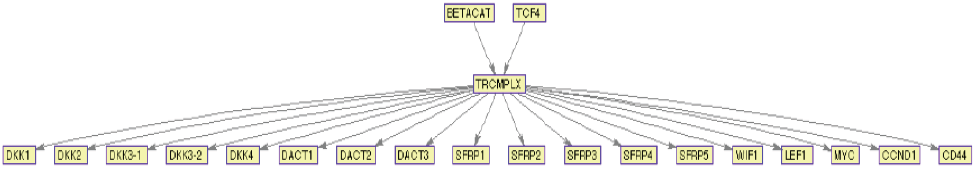
Influence diagram of 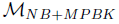 is a Naive Bayes model that contains minimal prior biological knowledge. In this model the state of *TRCMPLX* is assumed to be indicate whether the sample is cancerous or not.

~~~
% Generate dag for the interaction
% between nodeNames
dag = mk_adj_mat (interaction,…
 nodeNames, 0);
~~~

~~~
% To visualise the graphs or bayesian
% network
gObj = biograph (dag, nodeNames)
gObj = view (gObj);
~~~

Once the adjacency matrix is ready, the initialization of the Bayesian Network can be easily done. The total number of nodes is stored in `N` and the size of the nodes are defined in `nodeSizes`. In this project each node has a size of two as they contain discrete values representing binary states. Here the function ones defines a row vector with `N` columns. Thus each node is set to a size of 2. The total number of discrete nodes is defined in `discreteNodes`. Finally, the Bayesian Network is created using the function mk_bnet from the BNT that takes the following as input arguments (1) `dag` - the adjacency matrix (2) `nodeSizes` - defines the size of the nodes and (3) `discreteNodes` - the vector of nodes with their indices marked to be discrete in the Bayesian Network and dumps the network in the variable `bnet`.

~~~
% BN initialization
N = length (nodeNames); % # of nodes
~~~

~~~
% Define node sizes. NOTE - nodes are
% assumed to contain discrete values
nodeSizes = 2*ones (1, N);
~~~

~~~
% Discrete nodes
discreteNodes = 1:N;
~~~

~~~
% Create BN
bnet = mk_bnet (dag, nodeSizes,…
 ’names’, nodeNames, ’discrete’,…
 discreteNodes);
~~~

Section 4 of Sinha^1^ has been reproduced for completeness in tables 3, 4, 5, 6 and 7.

**Table 3.**
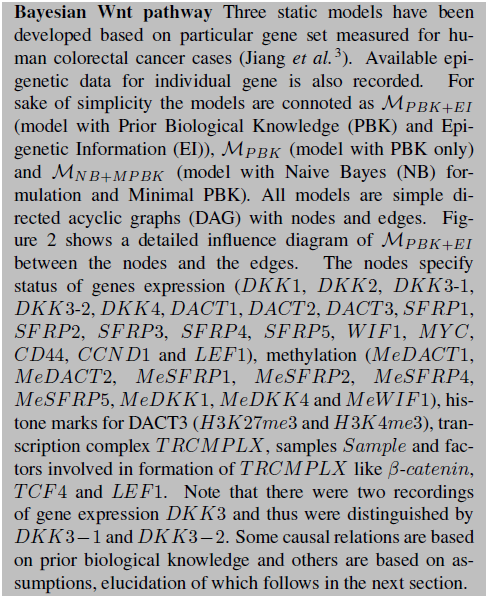
Bayesian Wnt pathway from Sinha ^1^

**Table 4.**
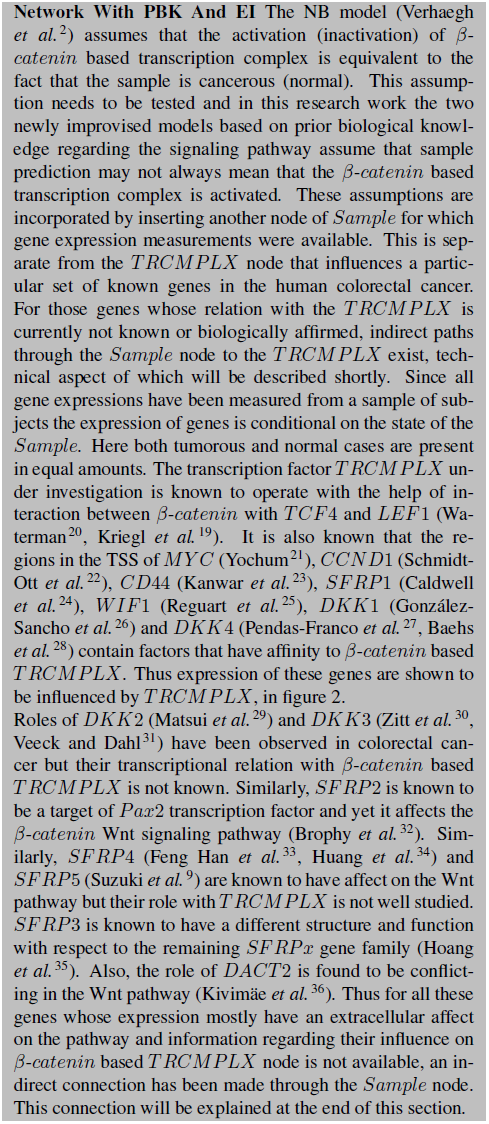
Network with PBK+EI from Sinha^1^

**Table 5.**
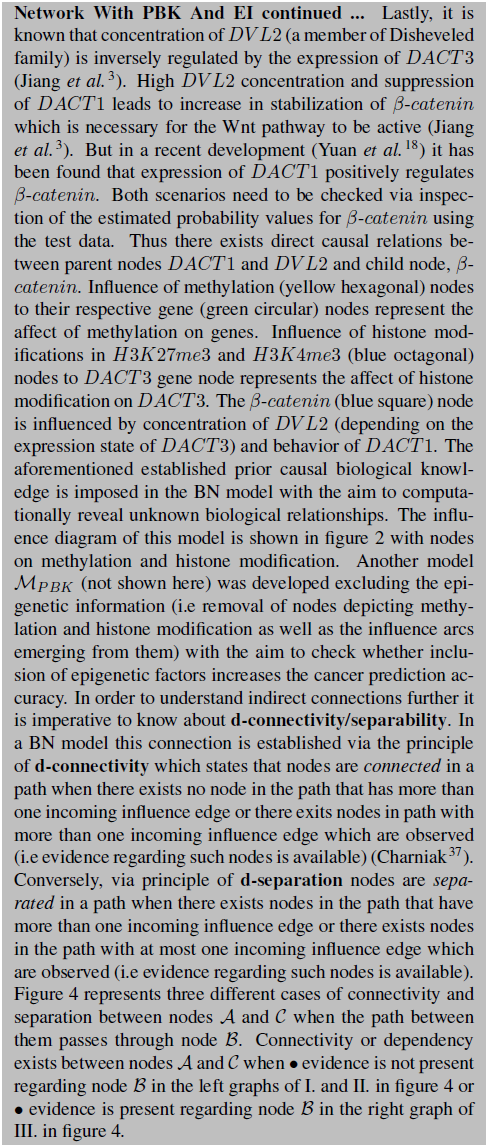
Network with PBK+EI continued from Sinha^1^

**Table 6.**
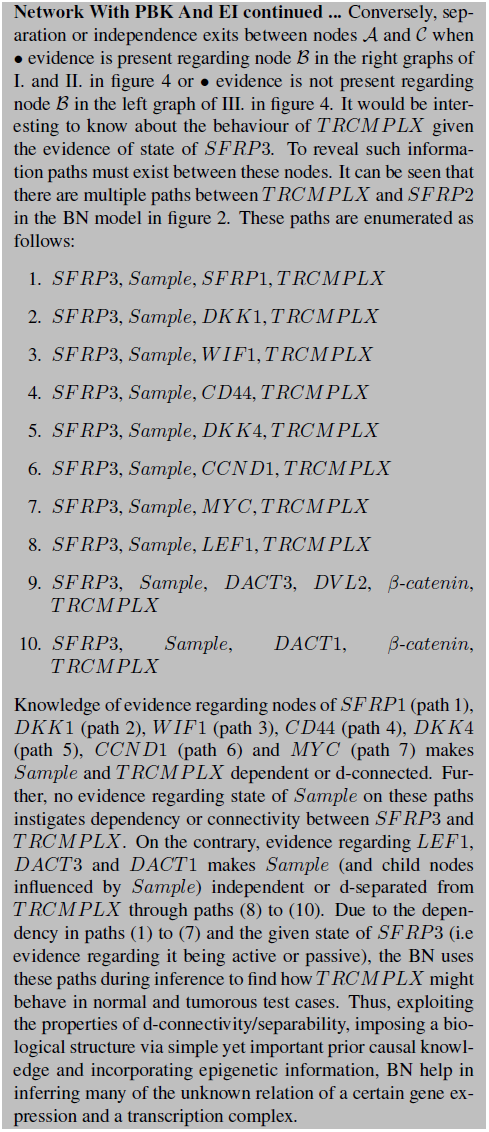
Network with PBK+EI continued from Sinha^1^

**Table 7.**
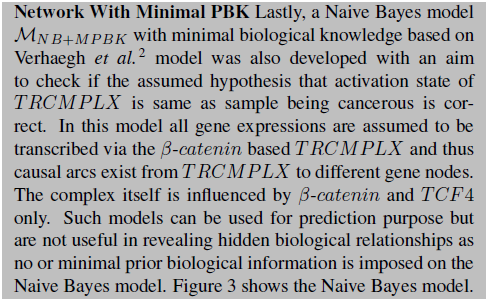
Network with NB+MPBK from Sinha ^1^

### 2.4 Hold out expriment

After the framework of the Bayesian Network has been constructed and initialized, the hold out experiment is conducted. The purpose of conducting the experiment is to generate results on different test data while training the Bayesian Network with different sets of training data, a multiple number of time. From Sinha^1^, the design of the experiment is a simple 2-holdout experiment where one sample from the normal and one sample from the tumorous are paired to form a test dataset. Excluding the pair formed in an iteration of 2-hold out experiment the remaining samples are considered for training of a BN model. Thus in a data set of 24 normal and 24 tumorous cases, a training set will contain 46 samples and a test set will contain 2 samples (one of normal and one of tumor). This procedure is repeated for every normal sample which is combined with each of the tumorous sample to form a series of test dataset. In total there will be 576 pairs of test data and 576 instances of training data. Note that for each test sample in a pair, the expression value for a gene is discretized using threshold computed for that particular gene from the training set. Computation of threshold will be elucidated later. This computation is repeated for all genes per test sample. Based on the available evidences from the state of expression of all genes, that constitute the test data, inference regarding the state of the both *β*-*catenin* transcription complex and the test sample is made. These inferences reveal • hidden biological relationship between the expressions of the set of genes under consideration and the *β*-*catenin* transcription complex and • information regarding the activation state of the *β*-*catenin* transcription complex and the state of the test sample, as a penultimate step to the proposed hypothesis testing. Two sample Kolmogorov-Smirnov (KS) test was employed to measure the statistical significance of the distribution of predictions of the states of the previously mentioned two factors.

Apart from testing the statistical significance between the states of factors, it was found that the prediction results for the factors, obtained from models including and excluding epigenetic information, were also significantly different. The receiver operator curve (ROC) graphs and their respective area under the curve (AUC) values indicate how the predictions on the test data behaved under different models. Ideally, high values of AUC and steepness in ROC curve indicate good quality results.

The hold out experiment begins with the computation of the total number of positive and negative labels present in the whole data set as well as the search of the indicies of the labels. For this the values in the variable `noSamples` and `transGroundTruthLabels` computed from function readCustomFile is used. `noPos` (`noNeg`) and `posLabelIdx` (`negLabelIdx`) store the number of positive (negative) labels and their indicies, respectively.

~~~
% Hold out experiment
%=====
% Compute no. of positive and negative
% labels and find indicies of both
noPos = 0;
posLabelIdx = [ ];
noNeg = 0;
negLabelIdx = [ ];
for i = 1:noSamples
      if transGroundTruthLabels(i) > 0
          noPos = noPos + 1;
           posLabelIdx = [posLabelIdx, i];
       else
          noNeg = noNeg + 1;
          negLabelIdx = [negLabelIdx, i];
      end
end
~~~

For storing results as well as the number of times the experiment will run, variables `runCnt` and `Runs` are initialized. The condition in the **if** statement is not useful now and will be described later.

~~~
runCnt = 0;
Runs = struct ([ ]);
if ~isempty (strfind (eviDence, ’me’))
      RunsOnObservedMethylation = …
        struct ([ ]);
end
~~~

For each and every positive (cancerous) and negative (normal) labels, the number of times the experiments runs is incremented in the count variable `runCnt`. Next the indicies for test data is separated by using the *i*^th^ positive and the *j*^th^ negative label and its indicies is stored in `testDataIdx`. The test data itself is then separated from `expressionMatrix` using the `testDataIdx` and stored in `dataForTesting`. The corresponding ground truth labels of the test data are extracted from `transGroundTruthLabels` using `testDataIdx` and stored in `labelForTesting`.

~~~
for i = 1:noPos
   for j = 1:noNeg
    % Count for number of runs
    runCnt = runCnt + 1;
    % Build test dataset (only 2
   % examples per test set)
   testDataIdx = [negLabelIdx(j),…
       posLabelIdx(i)];
   dataForTesting = expressionMatrix (:,…
         testDataIdx);
  labelForTesting = …
    transGroundTruthLabels (:,…
     testDataIdx);
~~~

After the storage of the test data and its respective indicies, `trainingDataIdx` is used to store the indicies of training data by eliminating the indicies of the test data. This is done using temporary variables `tmpPosLabelIdx` and `tmpNegLabelIdx`. `trainingDataIdx` is used to store the training data in variable `dataForTraining` using `expressionMatrix` and the indicies of training data in variable `labelForTraining` using `transGroundTruthLabels`.

~~~
% Remove test dataset from the whole
% dataset and build train dataset
tmpPosLabelIdx = posLabelIdx;
tmpNegLabelIdx = negLabelIdx;
tmpPosLabelIdx(i) = [ ];
tmpNegLabelIdx(j) = [ ];
trainDataIdx = [tmpNegLabelIdx,…
        tmpPosLabelIdx];
dataForTraining = …
  expressionMatrix (:,trainDataIdx);
labelForTraining = …
  transGroundTruthLabels (:,…
    trainDataIdx);
~~~

#### 2.4.1 Defining and estimating probabilities and conditional probabilities tables for nodes in bnet

Till now, the probabilities as well as conditional probability tables (cpt) for some of the nodes have been stored in the *probabilities.mat* file and loaded in the workspace. But the cpt for all the nodes in the `bnet` remain uninitialized. The next procedure is to initialize the tables using assumed values for some of the known nodes while estimating the entries of cpt for other nodes using training data.

To this end it is important to define a variable by the name `cpdStorage` of the format structure. Starting with all the nodes that have no parents and whose probabilities and cpt have been loaded in the workspace (saved in *probabilities.mat*), the **for** loop iterates through all the nodes in the network defined by `N`, stores the index of *k*^th^ node in `nodeidx` using function bnet.names with input argument `nodeNames`{`k`} and assigns values to cpt depending on the type of `model`. If 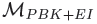 (model=’t1’) is used and the *k*^th^ entry in `nodeNames` matches with *TCF*4 then the cpt value in `PrTCF4` is assigned to `cpt`. The parent node of this node is assigned a value 0 and stored in `cpdStorage (k).parentnode`{`1`}. The name *TCF*4 or `nodeNames`{`k`} is assigned to `cpdStorage (k).node`. The cpt values in `cpt` is assigned to `cpdStorage (k).cpt`. Finally, the conditional probability density `cpt` for the node with name *TCF*4 is stored in `bnet.CPD` using function tabular CPD, the Bayesian Network `bnet`, the node index `nodeidx` and `cpt`. Similarly, values in `PrMeDKK1`, `avgPrMeDACT1`, `avgPrMeDACT2`, `avgPrH3K27me3`, `avgPrH3K4me3`, `PrMeSFRP1`, `PrMeSFRP2`, `PrMeSFRP4`, `PrMeSFRP5`, `PrMeWIF1` and `PrSample` initialize the cpt values for nodes *MeDACT*1, *MeDACT*2, *H*3*k*27*me*3, *H*3*k*4*me*3, *MeSF RP*1, *MeSF RP*2, *MeSF RP*4, *MeSF RP*5, *MeW IF*1 and *Sample*, respectively.

Similar initializations happen for models 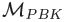 (model=’t2’) and 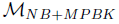 (model=’p1’). It should be noted that in 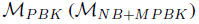 the only nodes without parents are *TCF* 4 and *Sample* (*TCF*4 and *BET ACAT*). To accomodate for these models, the necessary **elseif** statements have been embedded in the **for** loop below.

~~~
% Define P and CPD for the nodes of the
% bnet
cpdStorage = struct ([ ]);
% Store probabilities for nodes with no
% parents
for k = 1:N
  nodeidx = bnet.names (nodeNames{k});
  if isempty (bnet.parents{nodeidx})
  % tables for non-gene measurements
  if ~isempty (strfind (model,’t1’))
   if strcmp (nodeNames{k},’TCF4’)
    cpt = PrTCF4;
  elseif strcmp (nodeNames{k},…
    ’MeDKK1’) cpt = PrMeDKK1;
     elseif strcmp (nodeNames{k},…
    ’MeDACT1’)
  cpt = avgPrMeDACT1;
     elseif strcmp (nodeNames{k},…
  ’MeDACT2’)
    cpt = avgPrMeDACT2;
   elseif strcmp (nodeNames{k},…
  ’H3k27me3’)
  cpt = avgPrH3K27me3;
     elseif strcmp (nodeNames{k},…
  ’H3k4me3’)
  cpt = avgPrH3K4me3;
   elseif strcmp (nodeNames{k},…
  ’MeSFRP1’)
  cpt = PrMeSFRP1;
   elseif strcmp (nodeNames{k},…
  ’MeSFRP2’)
  cpt = PrMeSFRP2;
   elseif strcmp (nodeNames{k},…
  ’MeSFRP4’)
    cpt = PrMeSFRP4;
   elseif strcmp (nodeNames{k},…
  ’MeSFRP5’)
    cpt = PrMeSFRP5;
   elseif strcmp (nodeNames{k},…
  ’MeWIF1’)
  cpt = PrMeWIF1;
   elseif strcmp (nodeNames{k},…
  ’Sample’)
  cpt = PrSample;
   end
    elseif ~isempty (strfind (model,…
  ’t2’))
  if strcmp (nodeNames{k},’TCF4’)
    cpt = PrTCF4;
   elseif strcmp (nodeNames{k},…
    ’Sample’)
    cpt = PrSample;
     end
  elseif ~isempty (strfind (model,…
    ’p1’))
  if strcmp (nodeNames{k},’TCF4’)
    cpt = PrTCF4;
   elseif strcmp (nodeNames{k},…
    ’BETACAT’)
  cpt = PrBETACAT;
   end
 end
 cpdStorage (k) .parentnode {1} = 0;
 cpdStorage (k) .node = nodeNames {k};
 cpdStorage (k) .cpt = cpt;
 bnet .CPD {nodeidx} = tabular_CPD (…
  bnet, nodeidx, ’CPT’, cpt);
 end
end
~~~

In the same **for** loop above, the next step is to initialize probability as well as the cpt values for nodes with parents. Two cases exist in the current scenario, i.e nodes that (1) represent genes and (2) do not represent genes. To accomodate for gene/non-gene node classification a logical variable `GENE` is introduced. Also, before entering the second for loop described below, a variable `gene cpd` of the format structure is defined for storage of the to be computed cpt values for all genes in the data set. `parentidx` stores the index of the parents of the child node under consideration using the child’s index in `nodeidx` via `bnet.parents`{`nodeidx`}. The total number of parents a child node has is contained in `noParents`.

Initially `GENE` is assigned a value of 0 indicating that the node under consideration is not a gene node. If this is the case, the ~`GENE` in the **if** condition of the **for** loop below gets executed. In this case, depending on the type of the model cpt values of a particular node is initialized. For 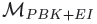 and 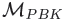 (model=’t1’ and model=’t2’), the cpt values for nodes *BETACAT*, *DV L*2 and *TRCMPLX* is stored using values in `PrBETACAT`, `PrDVL2` and `PrTRCMPLX`. As before, using function tabular_CPD and values in `nodeidx`, `bnet` and `cpt` as input arguments, the respective cpt is initialized in `bnet.CPD`{`nodeidx`}. Similar computations are done for 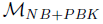 i.e model ’p1’ for node `TRCMPLX`. Finally, the indicies of the parents of the *k*^th^ child node is stored in `cpdStorage (k).parentnode`{`m`}.

On the other hand, if the name of the node in the *k*^th^ index of `nodeNames` matches the name in the index of `uniqueGenes`,a `parent` variable of format cell is defined within the second nested **for** loop below. The names of the parents are stored in this variable using `nodeNames`{`parentidx (n)`}. Next, the cpt values of these parent nodes are separately stored using a cell `parent cpd` and a count `cnt`. Finally, the cpd values for the *l*^th^ gene is determined using the function generateGenecpd in the script *generateGenecpd.m* that takes the following input arguments (1) `vecTraining` - gene expression of from training data (2) `labelTraining` - labels for training data (3) `nodeName` - name of the gene involved (4) `parent` name of parents of the child node or the gene under consideration (5) `parent cpd` - parent cpd values (6) `model` - kind of model and finally returns the output as a structure `gene cpd` containing cpd for the particular gene under consideration given its parents as well as a threshold value in the form of median. In the code below, the values of the following variables go as input arguments for the function generateGenecpd, in order (1) `dataForTraining (l,:)` - training data for the *l*^th^ unique gene, (2) `labelForTraining` - labels for training data, (3) `uniqueGenes`{`l`}, (4) `parent`, (5) `parent cpd`, (6) `model`. The output of the function is stored in the structure variable `x`. The threshold at which the probabilities were computed for the *l*^th^ gene is stored in `gene cpd (l).vecmedian` using `x.vecmedian` and the probabilities themselves are stored in `gene cpd (l).T` using `x.T`. These probabilities are reshaped into a row vector and stored in `cpt`. As mentioned before, using function tabular_CPD and values in `nodeidx`, `bnet` and `cpt` as input arguments, the respective cpt is initialized in `bnet.CPD`{`nodeidx`}. Finally, required values of cpt, name of *l*^th^ gene or *k*^th^ node and indicies of its parent nodes are stored in `cpdStorage (k).cpt, cpdStorage (k).node` and `cpdStorage (k) .parentnode`{`m`}, respectively.

It should be noted that the exposition of the generation of probability values for the different genes via the function generateGenecpd needs a separate treatment and will be addressed later. To maintain the continuity of the workflow of the program, the next step is addressed after the code below.

~~~
% Store probabilities for nodes with
% parents
gene_cpd = struct ([ ]);
for k = 1:N
  nodeidx = bnet .names (nodeNames {k});
  if ~isempty (bnet.parents {nodeidx})
   parentidx = bnet.parents {nodeidx};
   noParents = length (parentidx);
   GENE = 0;
   for l = 1:noGenes
     if strcmp (nodeNames {k},…
       uniqueGenes {l})
       % Find cpt of gene parent
       parent = { };
       for n = 1:noParents
         parent {n} = …
           nodeNames {parentidx (n)};
       end
       % Assign cpd to parent
       cnt = 0;
       parent_cpd = { };
       for m = 1:length (cpdStorage)
         for n = 1:noParents
           if strcmp (parent {n},…
             cpdStorage (m).node)
             cnt = cnt + 1;
             parent_cpd {cnt} = …
               cpdStorage (m).cpt;
           end
         end
       end
       x = generateGenecpd (…
        dataForTraining (l,:),…
        labelForTraining,…
        uniqueGenes {l}, parent,…
        parent_cpd, model);
      gene_cpd (l) .vecmedian = …
        x.vecmedian;
      gene_cpd (l).T = x.T;
      [r, c] = size (gene_cpd (l).T);
      cpt = reshape (gene_cpd (l).T,1,r*c);
      GENE = 1;
      break;
   end
end
% tables for non-gene measurements
if ~GENE
  if ~isempty (strfind (model,’t1’))
    if strcmp (nodeNames {k},’BETACAT’)
      cpt = PrBETACAT;
    elseif strcmp (nodeNames {k},’DVL2’)
      cpt = PrDVL2;
          elseif strcmp (nodeNames {k},’TRCMPLX’)
            cpt = PrTRCMPLX;
          end
         elseif ~isempty (strfind (model,’t2’))
           if strcmp (nodeNames {k},’BETACAT’)
            cpt = PrBETACAT;
          elseif strcmp (nodeNames {k},’DVL2’)
            cpt = PrDVL2;
           elseif strcmp (nodeNames {k},’TRCMPLX’)
             cpt = PrTRCMPLX;
           end
         elseif ~isempty (strfind (model,’p1’))
           if strcmp (nodeNames {k},’TRCMPLX’)
            cpt = PrTRCMPLX;
         end
       end
     end
     % record the parent index
     for m = 1:noParents
      cpdStorage (k).parentnode {m} = …
        parentidx (m);
     end
     cpdStorage (k).node = nodeNames {k};
     cpdStorage (k).cpt = cpt;
     bnet.CPD {nodeidx} = …
       tabular_CPD (bnet,nodeidx,’CPT’,cpt);
  end
end
~~~

#### 2.4.2 Evidence building and inference

The values estimated in `gene_cpd` as well as `cpdStorage` are stored for each and every run of the hold out experiment. Also, the dimensions of the testing data is stored.

~~~
% Function to store estimated
   % parameters
   Runs (runCnt) .geneCpd = gene_cpd;
   Runs (runCnt) .cpdStorage = cpdStorage;
~~~

~~~
   % Function to predict on test data
   % using trained BN
   [r, c] = size (dataForTesting);
~~~

Next, depending on the type of the evidence provided in `eviDence`, inferences can be made. Below, a section of code for evidence gene expression, which gets executed when the **case** ’ge’ matches with the parameter `eviDence` of the **switch** command, is explained. The issue that was to be investigated was whether the *β*-catenin based *TRCMPLX* is always switched on (off) or not when the *Sample* is cancerous (normal). In order to analyze this biological issue from a computational perspective, it would be necessary to observe the behaviour of the predicted states of both *TRCMPLX* as well as *Sample*, given all the available evidence. For this purpose, variable `tempTRCMPLXgivenAllge` is defined as a vector for each model separately, while variable `tempSAMPLE` is defined as a vector for biologically inspired models i.e 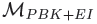 and 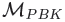 separately. This is so due to the assumption that the state of *TRCMPLX* is the same as the state of the test sample under consideration in the 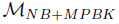 (a modification of Verhaegh *et al*.^2^).

In the section of the code below, **for** each of the test dataset an `evidence` variable of the format cell is defined. The `evidence` is of the size of equivalent to the number of node `N` in the network. Only those indicies in the cell will be filled for which information is available from the test data. Since the function twoHoldOutExp started with ’ge’ as an argument for type of evidence, `evidence` will be constructed from information available via gene expression from the test data. Thus for the *m*^th^ gene, if the gene expression in the test data (i.e `dataForTesting (m,k)`) is lower than the threshold generated using the median of expressions for this gene in the training data (i.e `gene cpd (m).vecmedian`), then the evidence for this gene is considered as inactive or repressed, i.e `evidence` {`bnet.names (uniqueGenes (m))`} = 1, else the evidence for this gene is considered active or expressed i.e `evidence` {`bnet.names (uniqueGenes (m))`} = 2. Iterating through all the genes, the `evidence` is initialized with the available information for the *k*^th^ test data.

Once the probability values have been initialized either by computation or assumption, then for the *k*^th^ test data, a Bayesian network engine is generated and stored in `bnetEngine` via the junction tree algorithm implemented in function `jtree_inf_engine` that uses the input argument as the newly initialized network stored in `bnet`. The `bnetEngine` is then fed with the values in `evidence` to generate a new engine that contains the updated probability values for nodes without evidence in the network. This is done using the function enter_evidence. According to BNT provided by Murphy *et al*.^15^, in the case of the jtree engine, enter_evidence implements a two-pass message-passing scheme. The first return argument (`engine`) contains the modified engine, which incorporates the evidence. The second return argument (`loglik`) contains the log-likelihood of the evidence. It is the first returned argument or the modified engine that will be of use further. It is important to note that for every iteration that points to a new test data in the **for** loop, a new Bayesian network engine is generated and stored in `bnetEngine`. If this is not done, then the phenomena of *explaining away* can occur on feeding new evidence to an already modified engine which incorporated the evidence from the previous test data. In *explaining away* the entrence of new evidence might out weigh the effect of an existing influencing factor or evidence thus making the old evidence redundant. This simulation is not related to such study of explaining away.

Finally, the belief that the *TRCMPLX* is switched on given the gene expression evidence i.e *Pr* (*TRCMPLX* = 2|ge as evidence) is computed by estimating the marginal probability values using the function marginal_nodes which takes the engine stored in `engine` and the name of the node using `bnet.names (’TRCMPLX’)`. The marginal probabilities are stored in `margTRCMPLX`. The final probability of *TRCMPLX* being switched on given all gene expression evidences is stored in `tempTRCMPLXgivenAllge` using `margTRCMPLX.T (2)`. Similarly, for biologically inspired models the belief that the test *Sample* is cancerous given the gene expression evidence i.e *Pr* (*Sample* = 2|ge as evidence) is computed using function marginal nodes that takes the engine stored in `engine` and the name of the node using `bnet.names` (’Sample’). The marginal probabilities are stored in `margSAMPLE`. The final probability of *Sample* being cancerous given all gene expression evidences is stored in `tempSAMPLE` using `margSAMPLE.T (2)`.

~~~
switch eviDence
  case ’ge’
    disp ([’Testing Example ’,…
      num2str (runCnt),…
      ’ – Based on all ge’]);
     tempTRCMPLXgivenAllge = [ ];
     if ~isempty (strfind (model, ’t’))
      tempSAMPLE = [ ];
     end
     % Build evidence for inference
     for k = 1:c
       evidence = cell (1,N);
     for m = 1:noGenes
     if dataForTesting (m,k) < = …
      gene_cpd (m) .vecmedian
      evidence {bnet.names (…
        uniqueGenes (m))} = 1;
     else
       evidence {bnet.names…
          (uniqueGenes (m))} = 2;
     end
     end
   % Build Bayesian engine
   bnetEngine = jtree_inf_engine (bnet);
   [engine, loglik] = …
   enter_evidence (bnetEngine,evidence);
   % Pr (TRCMPLX = 2|ge as evidence)
   margTRCMPLX = marginal_nodes (…
      engine, bnet.names (’TRCMPLX’));
   tempTRCMPLXgivenAllge = …
      [tempTRCMPLXgivenAllge,…
      margTRCMPLX.T (2)];
   if ~isempty (strfind (model, ’t’))
    % Pr (Sample = 2|ge as evidence)
    margSAMPLE = marginal_nodes (…
       engine,bnet.names (’Sample’));
    tempSAMPLE = [tempSAMPLE,…
       margSAMPLE.T (2)];
   end
end
~~~

Finally, for the particular count of the run of the experiment, `tempTRCMPLXgivenAllge` and `tempSAMPLE` are stored in the structure `Runs` using different variables associated with `Runs`. This iteration keeps happening until the two hold out experiment is exhausted. The case when `eviDence` is ’me’ or evidence for methylation will be discussed later as a programming project.

~~~
        % Function to store prediction values
        Runs (runCnt).condPrTRCMPLXgivenAllge…
          = tempTRCMPLXgivenAllge;
        if ~isempty (strfind (model,’t’))
          Runs (runCnt).condPrSAMPLE =…
          tempSAMPLE;
        end
      case ’me’
        % Project discussed later
    end
  end
end
~~~

### 2.5 Storing results, plotting graphs and saving files

The final section of the code deals with storing of the results, plotting of graphs and saving the results in the files. Since the current explanation is for gene expression evidence, the code pertaining to ’ge’ is explained. Readers might want to develop the code for evidence regarding methylation as a programming project.

To store results as well as the conditional probabilities for *TRCMPLX* and *SAMPLE* given all the gene expression evidence, a cell variable `Results`, a counter `cntResult` and vector variables `condPrTRCMPLXgivenAllge`, `condPrSAMPLE` and `labels` are defined as well as initialized. Next, the prediction values and original labels are stored while iterating through the total number of runs of the experiment. This is done using the **for** loop and the variable `runCnt`. For the *i*^th^ _run, predicted conditional probabilities of_ *TRCMPLX* and *Sample* from each run is stored in `condPrTRCMPLXgivenAllge(i,:)` and `condPrSAMPLE(i,:)`, depending on the model used. Finally, the ground truth labels of the test data are stored in a matrix were the *i*^th^ row is initialized with `labels(i,:) = [−1, +1];`. Here, labels it a matrix and −1 (+1) represent normal (cancerous) cases. Next, the variables `condPrTRCMPLXgivenAllge` and `condPrSAMPLE` are reshaped into vectors for further processing.

The plotting of the ROC curves and the estimation of their respective AUCs is achieved using function perfcurve that takes `labels`, either of the vectors `condPrTRCMPLXgivenAllge` or `condPrSAMPLE` depending on the type of model selected. The function churns out useful information in the form of the false positive rate in `X`, the true positive rate in `Y` and the estimated AUC for ROC of `condPrTRCMPLXgivenAllge` (`condPrSAMPLE`) in `AUCTRCMPLXgivenAllge` (`AUCSAMPLE`). The plot function is used to draw the graphs along with the depiction of legends using function legend. Finally, the two sample Kolmogorov-Smirnov test between the predictions of states of *TRCMPLX* and *Sample* is performed using the kstest2 function. This function takes the two vectors `condPrTRCMPLXgivenAllge` and `condPrSAMPLE` as arguments, compares the distribution of the predictions and returns the state of significance between the two distributions in `h01`. If the value of `h01` is 1, then statistical significance exists else it does not exist. Sinha^1^ shows that the statistical difference exists between predictions of *TRCMPLX* and *Sample* when the nodes for the same are segregated in the biologically inspired causal models, which is not the case with the naive Bayes model.

Lastly, the computed variables are stored in a .mat file using the function save. Options for using the save function can be obtained from the help command in Matlab.

~~~
if strcmp (eviDence, ’ge’)
  % Store results
  Results = { };
  cntResult = 0;
  % Estimation of performance levels
  condPrTRCMPLXgivenAllge = [ ];
  geneEvidence = { };
  if ~isempty (strfind (model, ’t’))
    condPrSAMPLE = [ ];
  end
  labels = [ ];
~~~

~~~
  % Store prediction values and
% original labels
for i = 1:runCnt
  condPrTRCMPLXgivenAllge(i,:) =…
    Runs(i) .condPrTRCMPLXgivenAllge;
geneEvidence {i} = Runs(i).geneEvidence;
  if ~isempty (strfind (model,’t’))
    condPrSAMPLE(i,:) =…
      Runs(i).condPrSAMPLE;
   end
   labels(i,:) = [−1, +1];
end
~~~

~~~
% Reshape the vectors
[r,c] = size (labels);
labels = reshape (labels,r*c,1);
condPrTRCMPLXgivenAllge =…
   reshape (condPrTRCMPLXgivenAllge,…
     r*c,1);
if ~isempty (strfind (model,’t’))
  condPrSAMPLE =…
    reshape (condPrSAMPLE, r*c,1);
end
~~~

~~~
% Plot the ROC curve and compute AUC
[X, Y, T, AUCTRCMPLXgivenAllge] =…
  perfcurve (labels,…
  condPrTRCMPLXgivenAllge,1);
plot (X, Y,’ r’);
xlabel (’False positive rate’);
ylabel (’True positive rate’);
if ~isempty (strfind (model,’t’))
  hold on;
  [X, Y, T, AUCSAMPLE] =…
   perfcurve (labels,condPrSAMPLE,1);
plot (X, Y,’ b’);
legend (’TRCMPLX – On’, ’SAMPLE – T’);
hold off;
~~~

~~~
% Perform ks–test the significance
% between models/evidences/predictions
[h01,p,ksstat] =…
  kstest2 (condPrTRCMPLXgivenAllge,…
  condPrSAMPLE);
end
~~~

~~~
if ~isempty (strfind (model,’t1’))
  save (’Results.mat’, ’Runs’,…
    ’condPrTRCMPLXgivenAllge’,…
    ’geneEvidence’, ’condPrSAMPLE’,…
    ’AUCTRCMPLXgivenAllge’,’AUCSAMPLE’,
    ’h01’);
elseif ~isempty (strfind (model,’ t2’))
  save (’Results.mat’,’Runs’,…
    ’condPrTRCMPLXgivenAllge’,…
    ’geneEvidence’,’condPrSAMPLE’,…
    ’AUCTRCMPLXgivenAllge’,’AUCSAMPLE’,
    ’h01’);
elseif ~isempty (strfind (model, ’p1’))
  save (’Results.mat’,’Runs’,…
    ’condPrTRCMPLXgivenAllge’,…
    ’geneEvidence’,…
    ’AUCTRCMPLXgivenAllge’);
  end
else
end
~~~

The ROC graphs and their respective AUC values found in the figures of Sinha^1^ are plotted by making variation in the assumed probability values of `PrTRCMPLX` in the function generateGenecpd that will be discussed later. Interpretations of the results can be studied in more depth from Sinha^1^.

Finally, a full section is dedicated to the computation of the probabilities for nodes with parents which has been implemented in function generateGenecpd.

### 2.6 Generating probabilities for gene nodes with parents

Here, the code for the function `generateGenecpd` is explained. As a recapitulation, the function generateGenecpd in the script *generateGenecpd.m* takes the following input arguments (1) `vecTraining` - gene expression of from training data (2) `labelTraining` - labels for training data (3) `nodeName` - name of the gene involved (4) `parent` - name of parents of the child node or the gene under consideration (5) `parent cpd` - parent cpd values (6) `model` - kind of model and finally returns the output as a structure `gene cpd` containing cpd for the particular gene under consideration given its parents as well as a threshold value in the form of median. In the code below, the values of the following variables go as input arguments for the function generateGenecpd, in order (1) `dataForTraining (l,:)` - training data for the *l*^th^ unique gene, (2) `labelForTraining` - labels for training data, (3) `uniqueGenes` {`l`}, (4) `parent`, (5) `parent cpd`, (6) `model`. The output of the function is stored in the structure variable `x`. The threshold at which the probabilities were computed for the *l*^th^ gene is stored in `gene cpd (l).vecmedian` using `x.vecmedian` and the probabilities themselves are stored in `gene cpd (l).T` using `x.T`.

The code begins with the storing of the dimension of a gene expression vector in `vecTraining` in `r` and `c` and recording the length of the vector containing the labels for the training data (in `labelTraining`) in `lencond`. Finally, the much reported threshold is estimated here using the median of the training data and stored in `vecmedian`.

~~~
% Rows is the gene expression and…
% columns are conditions (normal or
% cancerous)
[r, c] = size (vecTraining);
lencond = length (labelTraining);
~~~

~~~
% Take median as the threshold
vecmedian = median (vecTraining);
~~~

In Sinha^1^, the effect of *TRCMPLX* on the gene expression has been analysed as it is not known to what degree the *TRCMPLX* plays a role in the Wnt signaling pathway. To investigate this Sinha^1^ incorporated a parameter `p` that encodes the effect of *TRCMPLX* on the expression of the gene which is influenced by it. Thus while iterating through the list of parents if one encounters *TRCMPLX* as a parent, then `p` is initialized to a certain value. In Sinha^1^, the effect of *TRCMPLX* being active (1 − *p*) is incremented in steps of 0.1 from {0.5 to 0.9} and respective ROC graphs are plotted using the same.

~~~
% Defining affect of TRCMPLX on
% gene expression
noParents = length (parent);
for i = 1:noParents
  if ~isempty (strfind (model,’t’))
    if strfind (parent {i},’TRCMPLX’)
     p = 0.5;
    end
  end
end
~~~

It is important to note that the computation of gene probabilities differ from model to model and a detailed description of each computation is given for each gene for all three models, before explaining the computation for another gene. Also, from Sinha^1^, theoretically, for a gene *g_i_* ∀*i* genes, let there be *n_tr_* different instances of expression values from the sample training data. Let each of the *n_tr_* gene expression values be discretized to 0 and 1 based on their evaluation with respect to the median threshold. The 1’s represent the total number of expression where the gene is active and 0’s represent the total number of expression where the gene is inactive. In case of normal and tumorous samples, the proportions of 1’s and 0’s may be different. The median of the expression values is employed as a threshold to decide the frequency of *g_i_* being active or inactive given the state of the parent node (s). This median is also used along with the labels of the training data to decide the status of different parent factors affecting the gene under consideration.

#### 2.6.1 DKK1: (model=’t1’)

Since there are three parents for *DKK*1, namely *MeDKK*1, *Sample and TRCMPLX*, the cpt values for the table is segregated based on the status of methylation and quality of samples. A2 × 2 cross table for methylation and sample generates frequency estimates that can help derive probability values. The entries of the cross table depict the following cases (a) methylated in normal (represented by vector `mINn`) (b) un-methylated in normal (represented by vector `umINn`) (c) methylated in tumorous (represented by vector `mINt`) and (d) un-methylated in tumorous (represented by vector `umINt`), cases. For every *j^th^* entry in the `vecTraining`, if the label (`labelTraining(j)`) is normal (≤0) and the *DKK*1 gene expression (`vecTraining(j)`) is less than the estimated median (≤`vecmedian`) then value in `vecTraining(j)` is appended to `mINn`. Here, expression level lower than median indicates probable repression due to methylation in normal case. If the label (`labelTraining(j)`) is normal (≤0) and the *DKK*1 gene expression (`vecTraining(j)`) is greater than the estimated median (≥`vecmedian`) then value in `vecTraining(j)` is appended to `umINn`. Here, expression level greater than median indicates probable activation due to un-methylation in normal case. If the label (`labelTraining(j)`) is tumorous (≥0) and the *DKK*1 gene expression (`vecTraining(j)`) is less than the estimated median (≤`vecmedian`) then value in `vecTraining(j)` is appended to `mINt`. Here, expression level lower than median indicates probable repression due to methylation in tumorous case. And finally, If the label (`labelTraining(j)`) is tumorous (≥0) and the *DKK*1 gene expression (`vecTraining(j)`) is greater than the estimated median (≥`vecmedian`) then value in `vecTraining(j)` is appended to `umINt`. Here, expression level greater than median indicates probable activation due to un-methylation in tumorous case.

~~~
% Segregate values based on status
% of methylation and samples
mINn = [ ];
umINn = [ ];
mINt = [ ];
umINt = [ ];
for j = 1:lencond
  if labelTraining(j) < 0 && …
    vecTraining(j) < vecmedian
    mINn = [mINn, vecTraining(j)];
  elseif labelTraining(j) < 0 && …
    vecTraining(j) > = vecmedian
    umINn = [umINn, vecTraining(j)];
  elseif labelTraining(j) > 0 && …
    vecTraining(j) < vecmedian
    mINt = [mINt, vecTraining(j)];
  else
    umINt = [umINt, vecTraining(j)];
  end
end
~~~

Before estimating the values for cpt of *DKK*1, it is important to see how (1) the probability table would look like and (2) the probability table is stored in BNT (Murphy *et al.*^15^). Table 8 represents the conditions of sample as well as the methylation along with transcription complex and the probable beliefs of events (*DKK*1 being on/off). With three parents and binary state, the total number of conditions is 2^3^. To estimate the values of the probable beliefs of an event, the following computation is done. (**Case** - *TRCMPLX* is Off) The Pr (*DKK*1 On|*Sample -* Normal, *Me -* UM) being low, is the fraction of number of 1’s in the normal sample (`a`×*p*) and the sum of total number of normal samples and number of 1’s in the tumorous samples, i.e the non-methylated gene expression values in tumorous samples (`A`). Similarly, Pr (*DKK*1 - On|*Sample* Tumor,*Me -* UM) being low, is the fraction of number of 1’s in the tumorous sample (`b`×*p*) and the sum of total number of tumorous samples and number of 1’s in the normal samples, i.e the non-methylated gene expression values in normal samples (`B`). Again, Pr (*DKK*1 - Off|*Sample -* Normal,*Me* M) being high, is the fraction of number of 0’s in the normal sample (`c`×*p*) and the sum of total number of normal samples and number of 0’s in the tumorous samples, i.e the methylated gene expression values in tumorous samples (`C`). Finally, Pr (*DKK*1 - Off|*Sample -* Tumor,*Me -* M) being high, is the fraction of number of 0’s in the tumorous sample (`d`×*p*) and the sum of total number of tumorous samples and number of 0’s in the normal samples, i.e the methylated gene expression values in normal samples (`D`).

(**Case** - *TRCMPLX* is On) Next, the Pr (*DKK*1 On|*Sample -* Normal,*Me -* UM) being low, is the fraction of number of 1’s in the normal sample (`a`× (1 − *p*)) and the sum of total number of normal samples and number of 1’s in the tumorous samples, i.e the non-methylated gene expression values in tumorous samples (`A`). Similarly, Pr (*DKK*1 On|*Sample -* Tumor,*Me -* UM) being low, is the fraction of number of 1’s in the tumorous sample (`b`× (1−*p*)) and the sum of total number of tumorous samples and number of 1’s in the normal samples, i.e the non-methylated gene expression values in normal samples (`B`). Again, Pr (*DKK*1 - Off|*Sample* Normal,*Me -* M) being high, is the fraction of number of 0’s in the normal sample (`c`× (1−*p*)) and the sum of total number of normal samples and number of 0’s in the tumorous samples, i.e the methylated gene expression values in tumorous samples (`C`). Finally, Pr (*DKK*1 - Off|*Sample -* Tumor,*Me -* M) being high, is the fraction of number of 0’s in the tumorous sample (`d`× (1 −*p*)) and the sum of total number of tumorous samples and number of 0’s in the normal samples, i.e the methylated gene expression values in normal samples (`D`). Complementary conditional probability values for *DKK*1 being inactive

**Table 8.**
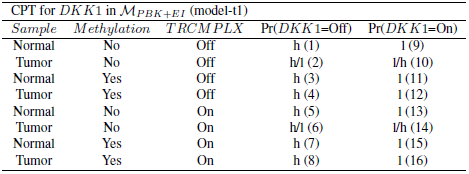
Conditional probability table for *DKK*1 in 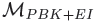 (model-t1). h - probability of event being high; l - probability of event being low. Serial numbers in brackets represent the ordering of numbers in vectorial format.

can easily be computed from the above estimated values.

~~~
% Generate frequencies for conditional
% probability values
~~~

~~~
% pr (DKK1 - On|Sample - Normal,Me - UM)
% # of On’s in Normal
a = length (umINn);
% total # of On’s in Normal and
% Unmethylation
A = length (umINn) + length (mINn)…
   + length (umINt);
~~~

~~~
% pr (DKK1 - On|Sample - Tumor,Me - UM)
% # of On’s in Tumor
b = length (umINt);
% total # of On’s in Normal and
% Unmethylation
B = length (umINt) + length (umINn)…
  + length (mINt);
~~~

~~~
% pr (DKK1 - Off|Sample - Normal,Me - M)
% # of Off’s in Normal
c = length (mINn);
% total # of Off’s in Normal and…
% Methylation
C = length (mINn) + length (umINn)…
   + length (mINt);
~~~

~~~
% pr (DKK1 - Off|Sample - Tumor,Me - M)
% # of Off’s in Normal
d = length (mINt);
% total # of Off’s in Normal and
% Methylation
D = length (mINt) + length (umINt)…
   + length (mINn);
~~~

These values are stored in variable `T` and the estimation is shown in the following section of the code. After the values in `T` has been established, a constant 1 is added as pseudo count to convert the distribution to a probability distribution via Dirichlet process. This is done to remove any deterministic 0/1 values appearing in the probability tables. If 0/1 appears in the probability tables then one has deterministic evidence regarding an event and the building of the Bayesian engine collapses. Finally, the frequencies in `T` are normalized in order to obtain the final conditional probability values for *DKK*1. Estimation of cpts for genes *SF RP* 1, *WIF* 1 and *DKK*4 which have methylation, *TRCMPLX* and *Sample* as parents require same computations as above. Figure 5 shows the pictorial representation of one of the cpt in 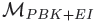

**Fig. 4.**
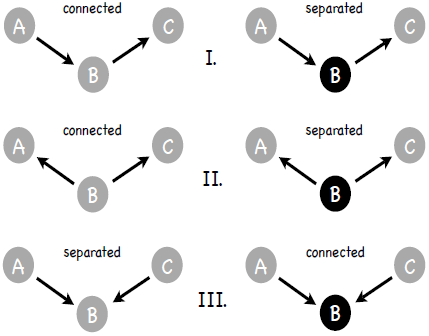
Cases for d-connectivity and d-separation. Black (Gray) circles mean evidence is available (not available) regarding a particular node.

**Fig. 5.**
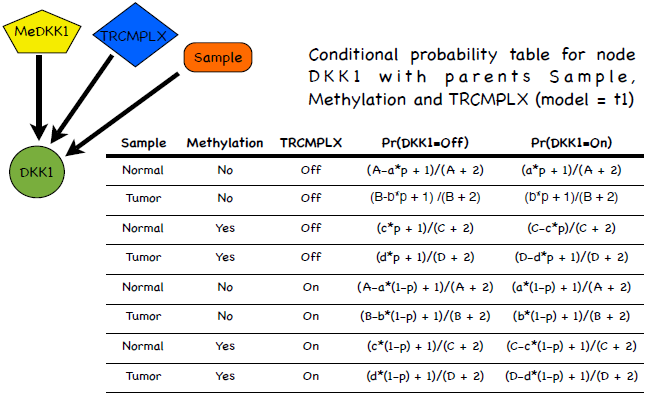
Conditional probability table for node *DKK*1 in 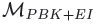.

~~~
% Multiply probability of TRCMPLX in
% on/off state to add the 3^rd^
% dimension in deciding the conditional
% probability tables.
~~~

~~~
% Conditional probability table for
% DKK1 given its parents
T = [A-a*p, a*p;…
  B-b*p, b*p;…
  c*p, C-c*p;…
  d*p, D-d*p;…
  A-a* (1-p), a* (1-p);…
  B-b* (1-p), b* (1-p);…
  c* (1-p), C-c* (1-p);…
  d* (1-p), D-d* (1-p)];
[r,c] = size (T);
% Convert the table to probability
% distribution via Dirichlet process
T = T + 1;
for i = 1:r
   T(i,:) = T(i,:)./sum (T(i,:));
end
~~~

(**model=’t2’**) There are two parents for *DKK*1, namely *TRCMPLX* and *Sample*. The conditional probability value for a gene being active or inactive is estimated based on the state of the *Sample*. But since the actual probability values for the activation of the *TRCMPLX* is not known the conditional probabilities are multiplied with a probability value of *p* when the *TRCMPLX* is off and with probability value 1−*p* when the *TRCMPLX* is on.

The analysis of quality of sample generates frequency estimates that can help derive probability values. These frequencies depict the following cases (a) gene repressed in normal (represented by vector `offINn`) (b) gene expressed in normal (represented by vector `onINn`) (c) gene repressed in tumorous (represented by vector `offINt`) and (d) gene expressed in tumorous (represented by vector `onINt`), cases. For every *j*^th^ entry in the `vecTraining`, if the label (`labelTraining(j)`) is normal (≤0) and the *DKK*1 gene expression (`vecTraining(j)`) is less than the estimated median (≤`vecmedian`) then value in `vecTraining(j)` is appended to `offINn`. Here, expression level lower than median indicates probable gene repression in normal case. If the label (`labelTraining(j)`) is normal (≤0) and the *DKK*1 gene expression (`vecTraining(j)`) is greater than the estimated median (≥`vecmedian`) then value in `vecTraining(j)` is appended to `onINn`. Here, expression level greater than median indicates probable gene activation in normal case. If the label (`labelTraining(j)`) is tumorous (≥0) and the *DKK*1 gene expression (`vecTraining(j)`) is less than the estimated median (≤`vecmedian`) then value in `vecTraining(j) is ap`pended to `offINt`. Here, expression level lower than median indicates probable gene repression in tumour case. And finally, If the label (`labelTraining(j)`) is tumorous (≥0) and the *DKK*1 gene expression (`vecTraining(j)`) is greater than the estimated median (≥`vecmedian`) then value in `vecTraining(j)` is appended to `onINt`. Here, expression level greater than median indicates probable gene activation in tumorous case.

~~~
% Segregate values based on
% status of TRCMPLX
onINn = [ ];
offINn = [ ];
onINt = [ ];
offINt = [ ];
for j = 1:lencond
  if labelTraining(j) < 0 &&…
    vecTraining(j) < vecmedian
    offINn = [offINn, vecTraining(j)];
  elseif labelTraining(j) < 0 &&…
    vecTraining(j) > = vecmedian
    onINn = [onINn, vecTraining(j)];
  elseif labelTraining(j) > 0 &&…
    vecTraining(j) < vecmedian
    offINt = [offINt, vecTraining(j)];
else
    onINt = [onINt, vecTraining(j)];
  end
end
~~~

Before estimating the values for cpt of *DKK*1, it is important to see how (1) the probability table would look like and (2) the probability table is stored in BNT (Murphy *et al.*^15^). Table 9 represents the conditions of *Sample* as well as *TRCMPLX* and the probable beliefs of events (*DKK*1 being on/off). With two parents and binary state, the total number of conditions is 2^2^. To estimate the values of the probable beliefs of an event, the following computation is done. The probability of gene expression being active given *Sample* is normal and *TRCMPLX* is off i.e Pr (*DKK*1 = Active |*Sample* = Normal, *TRCMPLX* = Off), is the fraction of number of 1’s in the normal sample (`a`×*p*) and the sum of total number of normal samples (`A`). Similarly, the probability of gene expression being active given *Sample* is tumorous and *TRCMPLX* is off i.e Pr (*DKK*1 = active |*Sample* = tumorous, *TRCMPLX* = Off), is the fraction of number of 1’s in the tumorous sample (`b`×*p*) and the sum of total number of tumorous samples (`B`). Again, the probability of gene expression being inactive given *Sample* is normal and *TRCMPLX* is on i.e Pr (*DKK*1 = inactive |*Sample* = normal, *TRCMPLX* = On), is the fraction of number of 0’s in the normal sample (`A-a`× (1 − *p*)) and the sum of total number of normal samples (`A`). Lastly, the probability of gene expression being inactive given *Sample* is tumorous and *TRCMPLX* is on i.e Pr (*DKK*1 = inactive |*Sample* = tumorous, *TRCMPLX* = On), is the fraction of number of 0’s in the tumorous sample (`B-b`× (1 − *p*)) and the sum of total number of tumorous samples (`b`). Complementary conditional probability values for *DKK*1 being inactive can easily be computed from the above estimated values.

**Table 9.** Conditional probability table for *DKK*1 in 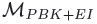 (model-t2). h - probability of event being high; l - probability of event being low. Serial numbers in brackets represent the ordering of numbers in vectorial format.

| CPT for *DKK1* in $\mathcal{M}_{PBK}$ (model-t2) | | | |
| --- | --- | --- | --- |
| *Sample* | *TRCMPLX* | $\Pr(DKK1=\text{Off})$ | $\Pr(DKK1=\text{On})$ |
| Normal | Off | h (1) | l (5) |
| Tumorous | Off | l (2) | h (6) |
| Normal | On | h (3) | l (7) |
| Tumorous | On | l (4) | h (8) |

~~~
% Generate frequencies for conditional
% probability values
% pr (DKK1 - On|Sample - N,TRCMPLX - Off)
% # of On’s when Sample is N
a = length (onINn);
% total # of TRCMPLX is Off
A = length (onINn) + length (offINn);
~~~

~~~
% pr (DKK1 - On|Sample - T,TRCMPLX - Off)
% # of On’s when Sample is T
b = length (onINt);
% total # of TRCMPLX is On
B = length (onINn) + length (offINt);
~~~

~~~
% Conditional probability table
% for DKK1 given its parents
T = [A-a*p, a*p;…
  B-b*p, b*p;…
  A-a* (1-p), a* (1-p);…
  B-b* (1-p), b* (1-p)];
[r,c] = size (T);
~~~

After the values in `T` has been established, a constant 1 is added as pseudo count to convert the distribution to a probability distribution via Dirichlet process. Finally, the frequencies in `T` are normalized in order to obtain the final conditional probability values for *DKK*1. Estimation of cpts for genes *SFRP* 1, *CCND*1, *CD*44, *WIF*1, *MYC* and *DKK*4 which has *TRCMPLX* and *Sample* as parents require same computations as above. Figure 6 shows the pictorial representation of one of the cpt in 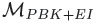

**Fig. 6.**
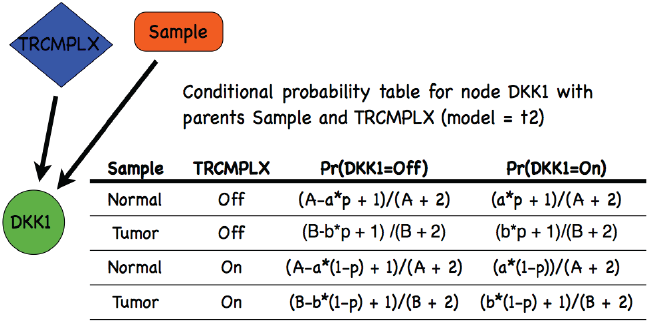
Conditional probability table for node *DKK*1 in 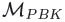.

~~~
% Convert the table to probability
% distribution via Dirichlet process
T = T + 1;
for i = 1:r
  T(i,:) = T(i,:)./sum (T(i,:));
end
~~~

(**model=’p1’**) Following the Naive Bayes model presented by Verhaegh *et al.*^2^ and making slight modifications to it, Sinha^1^ generated 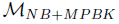. In this all genes have a single parent, namely *TRCMPLX* and it is assumed that the predicted state of *TRCMPLX* is exactly the same as the quality of the test sample. Thus the initial probability values for *TRCMPLX* are assumed to be fixed and no variation is made on it. The conditional probability value for a gene being active or inactive is estimated based on the state of the *TRCMPLX*.

The segregation of the probability values depends on the following conditions (a) gene is active and *TRCMPLX* is on (represented by vector `onINTrOn`) (b) gene is inactive and *TRCMPLX* is off (represented by vector `offINTrOn`) (c) gene is active and *TRCMPLX* is off (represented by vector `onINTrOff`) and (d) gene is inactive (represented by vector `offINTrOff`). For every *j*^th^ entry in the `vecTraining`, if the label (`labelTraining(j)`) is ≤0 (*TRCMPLX* is off) and the *DKK*1 gene expression (`vecTraining(j)`) is less than the estimated median (≤`vecmedian`) then value in `vecTraining(j)` is appended to `offINTrOff`. If the label (`labelTraining(j)`) is ≤0 (*TRCMPLX* is off) and the *DKK*1 gene expression (`vecTraining(j)`) is greater than the estimated median (≥`vecmedian`) then value in `vecTraining(j)` is appended to `onINTrOff`. If the label (`labelTraining(j)`) is ≥0 (*TRCMPLX* is on) and the *DKK*1 gene expression (`vecTraining(j)`) is less than the estimated median (≤`vecmedian`) then value in `vecTraining(j)` is appended to `offINTrOn`. And finally, if the label (`labelTraining(j)`) is ≥0 (*TRCMPLX* is on) and the *DKK*1 gene expression (`vecTraining(j)`) is greater than the estimated median (≥`vecmedian`) then value in `vecTraining(j) is ap`pended to `onINTrOn`.

~~~
% Segregate values based on
% status of TRCMPLX
onINTrOn = [ ];
offINTrOn = [ ];
onINTrOff = [ ];
offINTrOff = [ ];
for j = 1:lencond
  if labelTraining(j) < 0 &&…
    vecTraining(j) < vecmedian
    offINTrOff = [offINTrOff,…
    vecTraining(j)];
  elseif labelTraining(j) < 0 &&…
    vecTraining(j) >= vecmedian
    onINTrOff = [onINTrOff,…
    vecTraining(j)];
  elseif labelTraining(j) > 0 &&…
    vecTraining(j) < vecmedian
    offINTrOn = [offINTrOn,…
    vecTraining(j)];
  else
    onINTrOn = [onINTrOn,…
    vecTraining(j)];
end
~~~

Before estimating the values for cpt of *DKK*1, it is important to see how (1) the probability table would look like and (2) the probability table is stored in BNT (Murphy *et al.*^15^). Table 10 represents the conditions of *TRCMPLX* and the probable beliefs of events (*DKK*1 being on/off). With a single parent and binary state, the total number of conditions is 2^1^. To estimate the values of the probable beliefs of an event, the following computation is done. The probability of gene expression being active given *TRCMPLX* is off i.e Pr (*DKK*1 = Active |*TRCMPLX* = Off), is the fraction of number of 1’s in the normal sample (`a`) and the sum of total number of normal samples (`A`). Similarly, the probability of gene expression being inactive given *TRCMPLX* is off i.e Pr (*DKK*1 = active |*TRCMPLX* = On), is the fraction of number of 1’s in the tumorous sample (`b`) and the sum of total number of tumorous samples (`B`). Complementary conditional probability values for *DKK*1 being inactive can easily be computed from the above estimated values. Figure 6 shows the pictorial representation of one of the cpt in 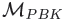

**Table 10.** Conditional probability table for *DKK*1 in 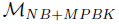 (model-p1). h - probability of event being high; l probability of event being low. Serial numbers in brackets represent the ordering of numbers in vectorial format.

| CPT for *DKK1* in $\mathcal{M}_{NB+PBK}$ (model-p1) | | |
| --- | --- | --- |
| *TRCMPLX* | $\Pr(DKK1=Off)$ | $\Pr(DKK1=On)$ |
| Off | h (1) | l (3) |
| On | h (2) | l (4) |

~~~
% Generate frequencies for
% conditional probability values
% pr (DKK1 - On | TRCMPLX - Off)
% # of On’s when TRCMPLX is Off
a = length (onINTrOff);
% total # of TRCMPLX is Off
A = length (onINTrOff) + length (offINTrOff);
~~~

~~~
% pr (DKK1 - On | TRCMPLX - On)
% # of On’s when TRCMPLX is On
b = length (onINTrOn);
% total # of TRCMPLX is On
B = length (onINTrOn) + length (offINTrOn);
~~~

~~~
% Conditional probability table
% for DKK1 given its parents
T = [A-a, a;…
  B-b, b];
[r,c] = size (T);
~~~

After the values in `T` has been established, a constant 1 is added as pseudo count to convert the distribution to a probability distribution via Dirichlet process. Finally, the frequencies in `T` are normalized in order to obtain the final conditional probability values for *DKK*1. Figure 7 shows the pictorial representation of one of the cpt in 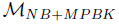

**Fig. 7.**
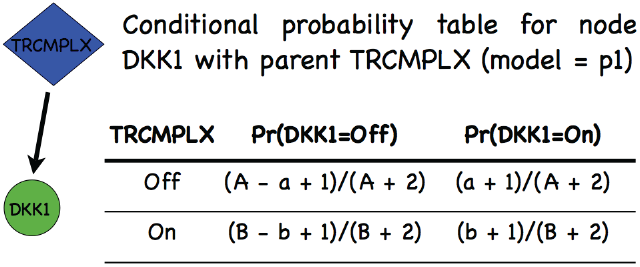
Conditional probability table for node *DKK*1 in 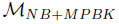.

~~~
% Convert the table to probability
% distribution via Dirichlet process
T = T + 1;
for i = 1:r
  T(i,:) = T(i,:)./sum (T(i,:));
end
~~~

#### 2.6.2 DKK2: (model-’t1’)

*Sample* is the single parent of *DKK*2. The conditional probability value for a gene being active or inactive is estimated based on the state of the *Sample*. The analysis of quality of sample generates frequency estimates that can help derive probability values. These frequencies depict the following cases (a) gene repressed in normal (represented by vector `offINn`) (b) gene expressed in normal (represented by vector `onINn`) (c) gene repressed in tumorous (represented by vector `offINt`) and (d) gene expressed in tumorous (represented by vector `onINt`), cases. For every *j*^th^ entry in the `vecTraining`, if the label (`labelTraining(j)`) is normal (≤0) and the *DKK*2 gene expression (`vecTraining(j)`) is less than the estimated median (≤`vecmedian`) then value in `vecTraining(j)` is appended to `offINn`. Here, expression level lower than median indicates probable gene repression in normal case. If the label (`labelTraining(j)`) is normal (≤0) and the *DKK*2 gene expression (`vecTraining(j)`) is greater than the estimated median (≥`vecmedian`) then value in `vecTraining(j)` is appended to `onINn`. Here, expression level greater than median indicates probable gene activation in normal case. If the label (`labelTraining(j)`) is tumorous (≥0) and the *DKK*2 gene expression (`vecTraining(j)`) is less than the estimated median (≤`vecmedian`) then value in `vecTraining(j)` is appended to `offINt`. Here, expression level lower than median indicates probable gene repression in tumour case. And finally, If the label (`labelTraining(j)`) is tumorous (≥0) and the *DKK*2 gene expression (`vecTraining(j)`) is greater than the estimated median (≥`vecmedian`) then value in `vecTraining(j)` is appended to `onINt`. Here, expression level greater than median indicates probable gene activation in tumorous case.

~~~
% Segregate values based on
% different types of samples
onINn = [ ];
offINn = [ ];
onINt = [ ];
offINt = [ ];
for j = 1:lencond
  if labelTraining(j) < 0 &&…
    vecTraining(j) < vecmedian
    offINn = [offINn, vecTraining(j)];
  elseif labelTraining(j) < 0 &&…
    vecTraining(j) > = vecmedian
    onINn = [onINn, vecTraining(j)];
  elseif labelTraining(j) > 0 &&…
    vecTraining(j) < vecmedian
    offINt = [offINt, vecTraining(j)];
  else
    onINt = [onINt, vecTraining(j)];
  end
end
~~~

Before estimating the values for cpt of *DKK*2, it is important to see how (1) the probability table would look like and (2) the probability table is stored in BNT (Murphy *et al*.^15^). Table 11 represents the conditions of *Sample and the prob*able beliefs of events (*DKK*2 being on/off). With a single parent and binary state, the total number of conditions is 2^1^. To estimate the values of the probable beliefs of an event, the following computation is done. The probability of gene expression being active given *Sample* is normal i.e Pr (*DKK*1 = Active |*Sample* = Normal), is the fraction of number of 1’s in the normal sample (`a`) and the sum of total number of normal samples (`A`). Similarly, the probability of gene expression being active given *Sample* is tumorous i.e Pr (*DKK*2 = active |*Sample* = Tumorous), is the fraction of number of 1’s in the tumorous sample (`b`) and the sum of total number of tumorous samples (`B`). Complementary conditional probability values for *DKK*2 being inactive can easily be computed from the above estimated values.

**Table 11.** Conditional probability table for *DKK*2 in 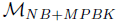 (model-t1). h - probability of event being high; l probability of event being low. Serial numbers in brackets represent the ordering of numbers in vectorial format.

| CPT for *DKK2* in $\mathcal{M}_{NB+PBK}$ (model-t1) | | |
| --- | --- | --- |
| *Sample* | Pr(*DKK2*=Off) | Pr(*DKK2*=On) |
| Normal | l/h (1) | h/l (3) |
| Tumor | h/l (2) | l/h (4) |

~~~
% Generate frequencies for
% conditional probability values
% pr (DKK2 - On | Sample - Normal)
% # of On’s in Normal
a = length (onINn);
% total # of samples in Normal
A = length (onINn) + length (offINn);
~~~

~~~
% pr (DKK2 - On | Sample - Tumor)
% # of On’s in Normal
b = length (onINt);
% total # of samples in Tumor
B = length (onINt) + length (offINt);
~~~

After the values in `T` has been established, a constant 1 is added as pseudo count to convert the distribution to a probability distribution via Dirichlet process. Finally, the frequencies in `T` are normalized in order to obtain the final conditional probability values for *DKK*2. Estimation of cpts for genes *DKK*3 − 1, *DKK*3 − 2, *SFRP*3 and *LEF*1 which have *Sample* as parent require same computations as above.

~~~
% Conditional probability table for
% DKK2 given its parents
T = [A-a, a;…
  B-b, b];
[r,c] = size (T);
~~~

~~~
% Convert the table to probability
% distribution via Dirichlet process
T = T + 1;
for i = 1:r
  T(i,:) = T(i,:)./sum (T(i,:));
end
~~~

(**model-’t2’**) When epigenetic factors are removed from 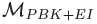 and the model transformed into 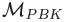 i.e model=’t2’, then the estimation of cpt values for *DKK*2 remain the same as in model=’t1’. Same computations apply for genes *DKK*3 − 1, *DKK*3 − 2, *SFRP*2, *SFRP*3, *SFRP*4, *SFRP*5, *LEF*1, *DACT*1, *DACT*2 and *DACT*3, in model=’t2’.

Figure 6 shows the pictorial representation of one of the cpt in 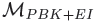 and 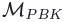

#### 2.6.3 DACT3: (model-’t1’)

The conditional probability value for a gene being active or inactive is estimated from generated frequency estimates that can help derive probability values. These frequencies depict the following cases (a) gene repressed in normal (represented by vector `offINn`) (b) gene expressed in normal (represented by vector `onINn`) (c) gene repressed in tumorous (represented by vector `offINt`) and (d) gene expressed in tumorous (represented by vector `onINt`), cases. For every *j*^th^ entry in the `vecTraining`, if the label (`labelTraining(j)`) is normal (≤0) and the *DACT*3 gene expression (`vecTraining(j)`) is less than the estimated median (≤`vecmedian`) then value in `vecTraining(j)` is appended to `offINn`. Here, expression level lower than median indicates probable gene repression in normal case. If the label (`labelTraining(j)`) is normal (≤0) and the *DACT* 3 gene expression (`vecTraining(j)`) is greater than the estimated median (≥`vecmedian`) then value in `vecTraining(j)` is appended to `onINn`. Here, expression level greater than median indicates probable gene activation in normal case. If the label (`labelTraining(j)`) is tumorous (≥0) and the *DACT* 3 gene expression (`vecTraining(j)`) is less than the estimated median (≤`vecmedian`) then value in `vecTraining(j) is ap`pended to `offINt`. Here, expression level lower than median indicates probable gene repression in tumour case. And finally, If the label (`labelTraining(j)`) is tumorous (≥0) and the *DACT* 3 gene expression (`vecTraining(j)`) is greater than the estimated median (≥`vecmedian`) then value in `vecTraining(j)` is appended to `onINt`. Here, expression level greater than median indicates probable gene activation in tumorous case.

~~~
% Segregate values based on status
% of histone repressive and active
% marks
onINn = [ ];
offINn = [ ];
onINt = [ ];
offINt = [ ];
~~~

~~~
for j = 1:lencond
    if labelTraining(j) < 0 &&…
      vecTraining(j) < vecmedian
      offINn = [offINn, vecTraining(j)];
    elseif labelTraining(j) < 0 &&…
      vecTraining(j) > = vecmedian
      onINn = [onINn, vecTraining(j)];
    elseif labelTraining(j) > 0 &&…
      vecTraining(j) < vecmedian
      onINt = [onINt, vecTraining(j)];
    else
      offINt = [offINt, vecTraining(j)];
  end
end
~~~

Before estimating the values for cpt of *DACT*3, it is important to see how (1) the probability table would look like and (2) the probability table is stored in BNT (Murphy *et al.*^15^). Table 12 represents the conditions of *Sample*, *H*3*K*4*me*3 and *H*3*K*4*me*3 the probable beliefs of events (*DACT*3 being on/off). Finally, from biological data presented in Jiang *et al.*^3^ the conditional probability values for the *DACT*3 gene being active based on the histone modification and the available samples suggest that *DACT*3 expression is high in normal samples when the histone repressive mark *H*3*K*27*me*3 is reduced and activating mark *H*3*K*4*me*3 are present in high abundance. Thus, the probability i.e Pr (*DACT*3 = *active*|*HK*327*me*3 = *low, H*3*K*4*me*3 = *high, Sample* = *normal*) is the fraction of the number of 1’s in the normal samples (`a`) and the total number of normal samples (`A`). For all other conditions of *H*3*K*27*me*3 and *H*3*K*4*me*3 when the *Sample* is normal the probability of *DACT* 3 being active is (`A-a`), i.e flip or complementray of Pr (*DACT* 3 = *active*|*HK*327*me*3 = *low, H*3*K*4*me*3 = *high, Sample* = *normal*). This is because in all other conditions of the histone marks the probability of *DACT* 3 being active will be reverse of what it is when *H*3*K*27*me*3 is reduced and *H*3*K*4*me*3 is present in abundance. Similarly, in case of tumorous samples, the probability of *DACT* 3 being active will occur when *H*3*K*27*me*3 is reduced and *H*3*K*4*me*3 is high abundance (a rare phenomena). Thus the probability i.e Pr (*DACT* 3 = *active*|*HK*327*me*3 = *low, H*3*K*4*me*3 = *high, Sample* = *tumorous*) is the fraction of the number of 1’s in the tumorous sample (`b`) and the total number of tumorous samples (`B`). For all other conditions of *H*3*K*27*me*3 and *H*3*K*4*me*3 when the *Sample* is tumorous the probability of *DACT* 3 being active is (`B-b`), i.e flip or complementray of Pr (*DACT* 3 = *active*|*HK*327*me*3 = *low, H*3*K*4*me*3 = *high, Sample* = *tumorous*). The reason for flip is the same as described above.

**Table 12.** Conditional probability table for *DACT*3 in 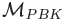 (model-t1). h - probability of event being high; l - probability of event being low. 1 - low; 2 - high.Serial numbers in brackets represent the ordering of numbers in vectorial format.

| CPT for *DACT3* in $\mathcal{M}_{PBK+EI}$ (model-t1) | | | | |
| --- | --- | --- | --- | --- |
| *H3K27me3* | *H3K4me3* | *Sample* | Pr(*DACT3*=Off) | Pr(*DACT3*=On) |
| 1 | 1 | Normal | h (1) | l (9) |
| 2 | 1 | Normal | h (2) | l (10) |
| 1 | 2 | Normal | l (3) | h (11) |
| 2 | 2 | Normal | h (4) | l (12) |
| 1 | 1 | Tumor | h (5) | l (13) |
| 2 | 1 | Tumor | h (6) | l (14) |
| 1 | 2 | Tumor | l (7) | h (15) |
| 2 | 2 | Tumor | h (8) | l (16) |

~~~
% Generate frequencies for
% conditional probability values
% pr (DACT3 - On | H3K27me3 - 1,
% H3K4me3 - 2, Sample - Normal)
% # of On’s in Normal
a = length (onINn);
% total # of On’s in Normal
A = length (offINn) + length (onINn);
~~~

~~~
% pr (DACT3 - On | H3K27me3 - 1,
% H3K4me3 - 2, Sample - Tumor)
% # of On’s in Tumor b = length (onINt);
% total # of On’s in Tumor
B = length (offINt) + length (onINt);
~~~

~~~
% In rest of the cases where
% (H3K27me3 - 1 and H3K4me3 - 2) is not
% present, the probabilities reverse.
~~~

After the values in `T` has been established, a constant 1 is added as pseudo count to convert the distribution to a probability distribution via Dirichlet process. Finally, the frequencies in `T` are normalized in order to obtain the final conditional probability values for *DACT* 3. Figure 9 shows the pictorial representation of one of the cpt in 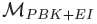

**Fig. 8.**
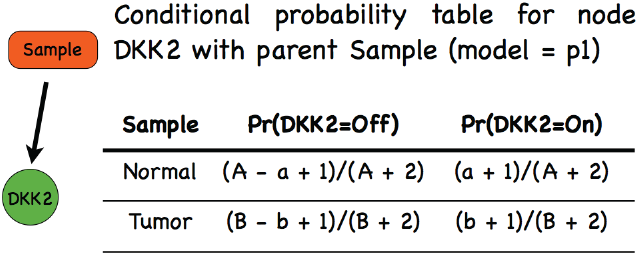
Conditional probability table for node *DKK*2 in 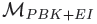 and 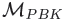

**Fig. 9.**
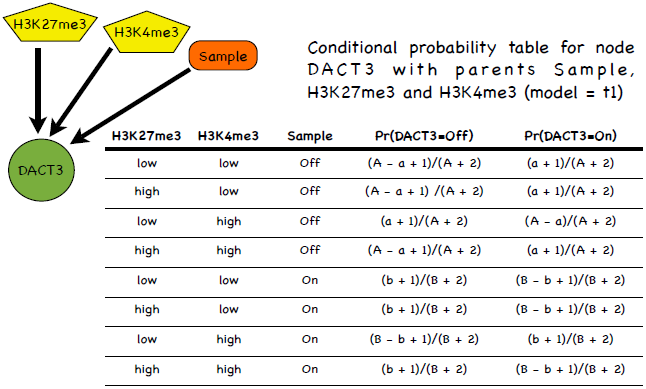
Conditional probability table for node *DACT*3 in 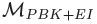.

~~~
% Conditional probability table
% for DACT3 given its parents
T = [a, A-a;…
   a, A-a;…
   A-a, a;…
   a, A-a;…
   b, B-b;…
   b, B-b;…
   B-b, b;…
   b, B-b];
[r,c] = size (T);
~~~

~~~
% Convert the table to probability
% distribution via Dirichlet process
T = T + 1;
for i = 1:r
  T(i,:) = T(i,:)./sum (T(i,:));
end
~~~

Finally, for every gene, after the computation of the probability values in their respective cpt, the function generate-Genecpd returns the following arguments as output.

~~~
gene_cpd = struct ();
gene_cpd.vecmedian = vecmedian;
gene_cpd.T = T;
~~~

## 3 A programming project for practice

To get a feel of the project, interested readers might want to implement the following steps when the evidence `eviDence` is ’me’. The code needs to be embedded as a case in the **switch** part of the twoHoldOutExp function. The idea is to perturb the methylation nodes with binary values and find if one can converge to the correct prediction of state of *TRCMPLX* as well as the *Sample*. These binary values are stored in a vector and represents a permutation of the methylation states of the methylation node in 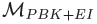. Varying the values of the vector can help study how perturbations affect the prediction of the network and the predictions. The steps are given below -

1. Define variables for storing predictions of *TRCMPLX* (`tempTRCMPLX`) and *Sample* (`tempSample`).
2. Find the total number of methylation cases in 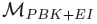 and store the number in a variable `noMethylation`.
3. Generate binary values for `noMethylation` nodes. Define a cell (`binaryStatesOfMethylation`) that can store vectors of binary values where every permutation represents a set of methylation states. The total number of permutations should be 2*^noMethylation^* which is stored in `noMethylationConfig`. One might want to use quantizer and num2bin functions from matlab.
4. Next, generate methylation evidences. Define a 2D matrix variable `methylationEvidence` that stores the methylation evidences. One might want to use the mat-lab function str2num. Finally, add a value of 1 to methylationEvidence as the BNT takes in ’1’ and ’2’ as states representing binary values.
5. Build evidence for inference for every test example. The steps following might be necessary

- For every methylation configuration and for every methylation node build evidence.
- Build a new bayesian network in `bnetEngine` using `jtree inf engine` and store the modified engine (in `engine`) using the function `enter evidence`.
- Finally, compute the Pr (*TRCMPLX* = 2|ge as evidence) and Pr (*Sample* = 2|ge as evidence) using the function marginal nodes.
6. Store predicted results on observed methylation in structure `Runs` indexed with `runCnt`.

After the section of new code is filled in, run the code and check the results.

## 4 Conclusion

A pedagogical walkthrough of a computational modeling and simulation project is presented using parts of programming code interleaved with theory. The purpose behind this endeavour is to acclimatize and ease the understanding of beginner students and researchers in transition, who intend to work on computational signaling biology projects. To this end, static Bayesian network models for the Wnt signaling pathway has been selected for elucidation. This is done due to lack or paucity of manuscripts explaining the computational experiments from tutorial perspective due to restrictive policies.

## Acknowledgement

Thanks to - (1) All anonymous reviewers who have helped in refining this manuscript. (2) Netherlands Bioinformatics Centre (NBIC) for funding the project. (3) Dr. ir. R. H. J. Fastenau (Dean of Faculty of Electrical Engineering Mathematics and Computer Science), Dr. Prof. Ir. K. Ch. A. M. Luyben (Rector Magnificus) and Drs. D. J. van den Berg (President) at Delft University of Technology, for providing support for conducting this work and giving permission to submit the manuscript. (4) Dr. Wim Verhaegh (faculty at Netherlands Bioinformatics Centre and a principal scientist at Molecular Diagnostic Lab in Philips Research) for providing technical details of Naive Bayes model for replicating experiments in Verhaegh *et al.*^2^. (5) Dr. Marcel J. T. Reinders (scientific director of Bioinformatics Research at Netherlands Bioinformatics Centre and a professor at Delft University of Technology) for refining the proposed model and suggesting inclusion of methylation data. (6) Dr. Robert P.W. Duin (retired associate professor at Delft University of Technology) for informal discussions on 2-holdout experiments and (7) Dr. Jeroen de Ridder (assistant professor at Delft University of Technology) and (8) Phd candidate Marc Hulsman (Delft University of Technology), for informal discussions on estimation of probability values on methylation data. (9) The author is indebted to Mr. Prabhat Sinha and Mrs. Rita Sinha for financially supporting this project while the author was on educational and work leave.

